# Mitigating nitrous oxide emission by an ultra-fast bioprocess enabling the removal of high concentration N_2_O

**DOI:** 10.1101/2024.10.08.615939

**Authors:** Ryota Maeda, Mikiko Sato, Kiwamu Minamisawa, Kengo Kubota

## Abstract

Nitrous oxide (N₂O) is known as a greenhouse gas as well as an ozone-depleting substance. Wastewater treatment process is one of the sources of N_2_O emission, and the high concentrations of N₂O in off-gas were reported from an anaerobic ammonium oxidation process. This study developed a novel N₂O removal process using a down-flow hanging sponge reactor to remove high concentrations of N₂O. More than 96% removal efficiencies were achieved for up to 300 ppm N₂O with 3 min gas retention time (GRT), and more than 99% removal efficiency was obtained for 2,000 ppm N_2_O with 18 min GRT. A maximum removal rate of 161 ± 26 mg-N/L-reactor/day was achieved, that was over 10 times faster than the pioneering process. Kinetic analysis indicated that the N₂O dissolution rate is a crucial factor in determining the N₂O removal rate in the reactor. Various N₂O reducers belonging to both clade I and II were detected in reactors, and *Azonexus* was thought to play a key role.

## 1. Introduction

Nitrous oxide (N_2_O) is a greenhouse gas and an ozone-depleting substance (Ravishankara et al., 2009). Its atmospheric concentration has increased significantly over the last 100 years, reaching 336 ppb in 2022 (IPCC, 2022; Tian et al., 2020). N_2_O has approximately 273 times greater global warming potential than that CO_2_ and has a lifetime of 114 years in the atmosphere. The recent increase in N_2_O emissions exceeds the worst-emission scenario of the IPCC; therefore, the reduction of N_2_O emissions is an urgent issue globally (Tian et al., 2020). The agricultural sector is the main source of N_2_O emissions, accounting for more than half of human activity-related emissions. Although their contribution is much lower, the waste and wastewater treatment sectors are also sources of N_2_O.

In wastewater treatment, N_2_O is generated in both aerobic and anaerobic processes related to nitrogen removal (i.e. nitrification and denitrification) by biotic and abiotic transformation processes (e.g. nitrifier denitrification, hydroxylamine oxidation, and incomplete denitrification) (Hallin et al., 2018; Sabba et al., 2018). Earlier studies identified several factors that influence N_2_O generation from wastewater treatment processes (e.g. dissolved oxygen (DO), nitrite and ammonium concentrations, and the chemical oxygen demand (COD) /N ratio) (Domingo-Félez and Smets, 2019; Kampschreur et al. 2009; Vasilaki et al. 2019; Wu et al. 2020). N_2_O emission mitigation approaches, such as DO control (Duan et al., 2021) and COD/N ratio control through carbon source input (Hu et al., 2013; Peng et al., 2017) were reported. However, these approaches are not necessarily universal because they are only effective under specific conditions (Han et al., 2023; Oba et al., 2024). As an alternative approach, processes removing N_2_O in off-gas using microbial reactions were developed (e.g., utilising a fixed bed reactor packed with Kaldnes rings (Frutos et al., 2016) or biofilters filled with polyurethane foam (Han et al., 2023; Yoon et al., 2019, 2017)). These studies targeted 100–300 ppm of N_2_O in either N_2_ (anaerobic) or air (aerobic) and successfully reduced N_2_O in the gaseous phase.

In addition to conventional nitrification-denitrification processes, anaerobic ammonium oxidation (anammox) is being considered for the removal of nitrogen from wastewater. More than 100 anammox-based full-scale plants are used globally to treat industrial and municipal wastewater (Lackner et al., 2014). The generation of N_2_O from anoxic anammox tanks were reported (Ali et al., 2016; Kampschreur et al., 2009, 2008; Okabe et al., 2011; Vasilaki et al., 2019) with concentrations reaching 1,300 ppm (laboratory-scale) (Okabe et al., 2011) and 4,000 ppm (full-scale) (Kampschreur et al., 2008) in off-gas, which is much higher than that of the previously tested concentrations for N_2_O removal (200 ppm in N_2_) (Yoon et al., 2017).

To remove high concentrations of N_2_O generated from anaerobic/anoxic wastewater treatment processes, such as anammox processes, we focused on down-flow hanging sponge (DHS) processes. DHS was originally developed for wastewater treatment, maintaining high sludge biomass in sponge media and resulting in effluent water quality comparable to that of conventional activated sludge processes. It allows aerobic treatment without aeration by supplying oxygen from the air due to gas-liquid equilibrium, while wastewater flows from top to bottom due to gravity (Kubota et al., 2024). The application of DHS reactors for gaseous substances such as hydrogen sulfide and toluene were reported by using the closed reactors (Tanikawa et al., 2022; Yamaguchi et al., 2018).

In this study, we developed a process that rapidly removed high concentrations of N_2_O using DHS. The N_2_O removal performance of the DHS reactors was evaluated by feeding different concentrations of N_2_O (5–2,000ppm) and supplying the supernatant from an anaerobic sewage sludge digester as a carbon source. Microelectrode experiments were conducted to assess the kinetics of the N_2_O reduction. Eventually, the microorganisms that may be involved in N_2_O removal were identified using 16S rRNA gene amplicon and shotgun metagenome analyses.

## 2. Material and Methods

### 2.1 Reactor configuration and operation

A DHS reactor consisting of fifteen sponge carriers with polyethene plastic net rings was placed in a closed column with a volume of 1.23 L. The carriers were vertically connected and suspended in the column. The size of polyurethane sponge was 30 mm in diameter and 30 mm in height. The total sponge volume was 0.32 L, resulting in a packing density of 26%.

Anaerobic sludge obtained from an anaerobic digester treating sewage sludge under mesophilic conditions was used as the seed sludge. The supernatant of the digested sludge was used as a carbon source and fed from the top of the reactor with a hydraulic retention time (HRT) of 3–24 hours based on the sponge volume. Feeding was performed intermittently using a timer. N_2_ gas containing 5, 40, 300 or 2,000 ppm N_2_O was subjected to the closed column from the inlet near the bottom of the reactor. The outlet for the treated gas was placed near the top of the reactor, resulting in gas flow from the bottom to the top. The gas flow rate was adjusted using a gas meter (RK1200; KOFLOC, Kyoto, Japan). The gas retention time (GRT) was adjusted by varying the gas flow rate. The reactor was set in an incubation chamber and operated anaerobically at 25°C in the dark. The GRT was calculated by using a gas volume at 25°C and an effective reaction volume of 0.91 L. Nitrogen loading of the reactor was calculated by using a reactor volume of 1.23 L and the amount of nitrogen under the standard temperature (0°C). A schematic of the experiment is shown in Fig.1.

**Fig. 1.**
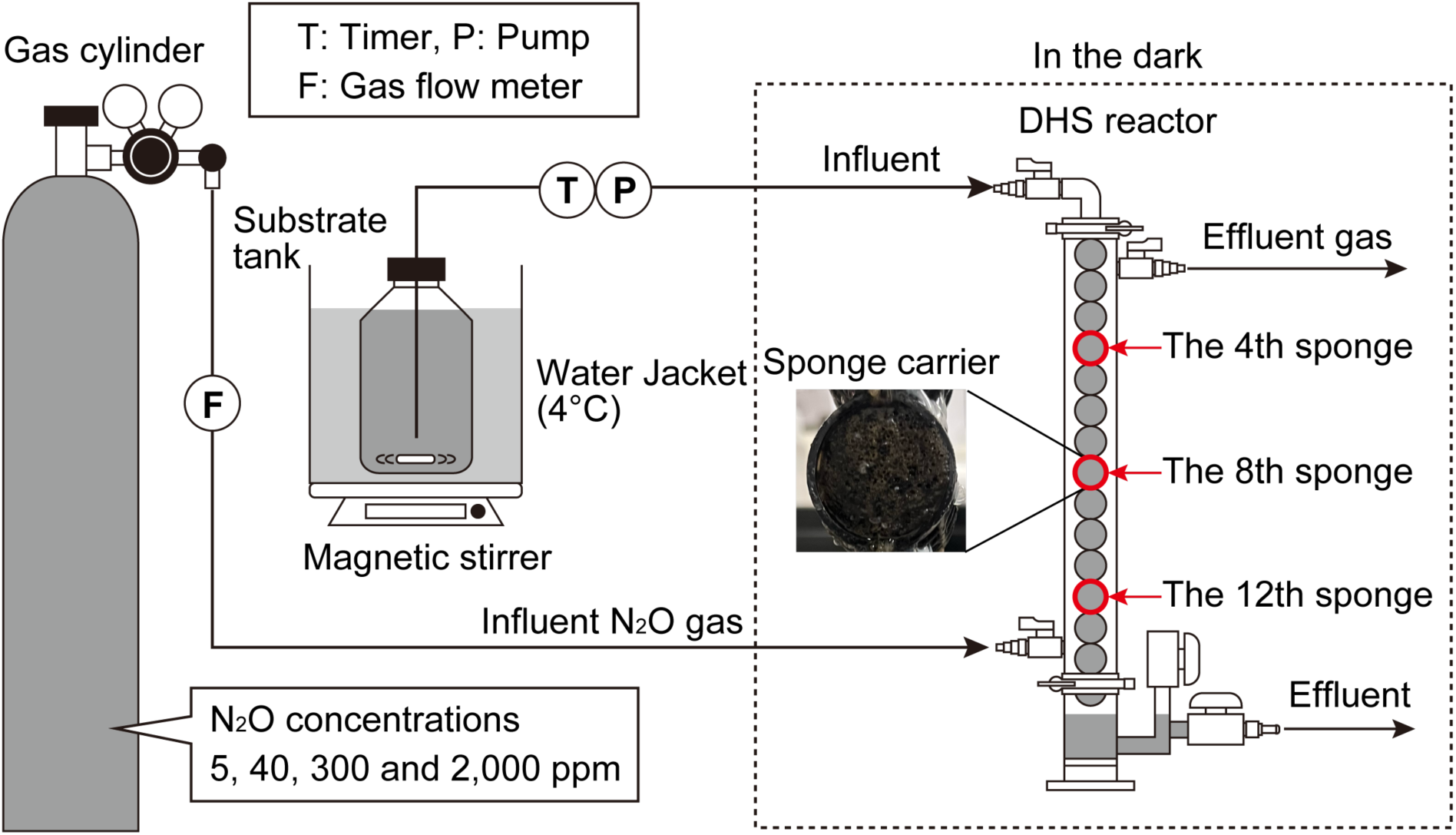
Schematic diagram of the experiment. The sludge samples collected from the 4th, 8th and 12th sponges from the top were used for the microelectrode experiments. The 4th and 12th sponges were used for 16S rRNA amplicon and metagenome analyses.

### 2.2 Measurement of gaseous N_2_O concentration

The effluent gas from the closed DHS reactor was measured every 1–2 days to evaluate N_2_O removal performance. In this study, two sets of electron capture detector gas chromatography (ECD-GC) (GC-2014, Shimadzu, Kyoto, Japan) with different columns were used, depending on the N_2_O concentration. Both ECD-GCs were equipped with a 63Ni ECD, and N_2_ was used as the carrier gas. For the measurement of lower N_2_O concentrations (< approximately 40 ppm), an ECD-GC with a tandem Porapak Q column (80/100 mesh; 3.0 mm × 1.0 m and 3.0 mm × 2.0 m, GL Sciences, Japan) was used. The temperatures were set at 80°C, 100°C, and 340°C for the column, injector, and detector, respectively. For the measurement of higher N_2_O concentrations (> approximately 40 ppm), an ECD-GC with a TC-BOND Q column (0.53 mm × 30 m, df=20 μm, GL Sciences, Japan) was used. The temperatures were set at 50°C, 280°C, and 300°C for the column, injector, and detector, respectively.

### 2.3 Water quality analysis and sludge sampling

Influent water and effluent water from the DHS reactor were sampled to evaluate changes in water quality. COD was measured using a HACH water quality analyser (DR890, HACH, Loveland, CO, USA).

Sludge samples were collected on the final experimental day. A sponge was immersed in 0.05 × phosphate buffered saline (PBS; 130 mM NaCl, 10.8 mM Na_2_HPO_4_, 4.2 mM NaH_2_PO_4_, pH 7.4) in a plastic bag, and sludge was suspended in PBS by squeezing the sponges. Part of the suspended sludge was used for microelectrode experiments (see Section 2.4). The other part was centrifuged at 2935×g for 15 min at 4°C. After removing the supernatant, a portion of the sludge was stored at -20°C for DNA extraction. The other part was used to measure the suspended solids (SS) and volatile suspended solids (VSS) according to the standard methods (APHA, 2017).

### 2.4 Microelectrode experiment for measuring N_2_O consumption rate

The N_2_O consumption rates of the sludge samples were measured using an N_2_O micro-respiration system with amperometric microsensors (Unisense, Aarhus, Denmark) (Fig. S1). Measurements and subsequent data analyses were conducted in accordance with an earlier study (Suenaga et al., 2018). Briefly, the sludge samples collected from the 4th, 8th and 12th sponges (Fig. 1) were used for the experiments. The sludge samples were initially diluted with anaerobic sludge digester supernatant, which was filter sterilised by using 0.22 µm Stericup^®^ filter (S2GVU01RE, Merck Millipore, Germany) to make sludge concentration suitable for the microelectrode experiment. Approximately 8 ml double-port chamber (Unisense, Aarhus, Denmark) was filled with the sample and placed in a water bath controlled at 25°C. N_2_O sensors were inserted into the chamber, and a certain amount (30–40 µl) of N_2_O water prepared by aerating pure N_2_O gas was injected into the chamber using a Hamilton syringe. The sample was stirred at 600 rpm, and the N_2_O concentration was continuously recorded using SensorTrace Suite ver. 2.8.0 (Unisense). After the injected N_2_O was completely consumed, fresh N_2_O water was injected, and N_2_O consumption was measured.

For the data analysis, noise removal and plot smoothing were performed using Sigma Plot 13.0 (Systat Software, San Jose, CA, USA). The theoretical maximum consumption rate of N_2_O (*V_max_*) was obtained using the Michaelis-Menten equation and the Excel solver function (Suenaga et al., 2018). The theoretical maximum N_2_O removal rate per reactor volume (mg-N/L-reactor/day) was calculated by multiplying *V_max_* (N_2_O removal rate per VSS) by the amount of VSS in the reactor.

### 2.5 16S rRNA gene amplicon sequence analysis

Sludge samples collected from the 4th and 12th sponges of each reactor were subjected to 16S rRNA gene amplicon sequencing analysis. DNA extraction was performed using ISOIL for Beads Beating kits (Nippon Gene, Tokyo, Japan), according to the manufacturer’s instructions. Prokaryotic 16S rRNA genes were amplified using 341f (5’-CCTAYGGGGRBGCASCAG-3’) and 806r mix (806r (5’-GGACTACHVGGGGTHTCTAAT-3’) and 806r-P (GGACTACCAGGGTATCTAAG-3’) in a 30:1 ratio) (Matsubayashi et al., 2017). The PCR conditions for library preparation were as described earlier (Ni et al., 2020). 16S rRNA gene amplicon sequencing using the MiSeq Reagent Kit v3 600 cycles was outsourced to the Bioengineering Lab. Co., Ltd. (Kanagawa, Japan). Nucleotide sequence data are available in the DDBJ Sequence Read Archive accession number PRJDB18886.

Data analysis was conducted using QIIME2 ver. 2023.7 (Bolyen et al., 2019). Quality trimming, primer sequence removal, paired-end assembly, and chimaera checking of the raw 16S rRNA gene sequencing data were performed using DADA2 (Callahan et al., 2016). The 16S rRNA gene sequences with ≥ 97% identity were clustered into operational taxonomic units (OTUs) using VSEAECH (Rognes et al., 2016). The OTUs were classified using the Greengens2 database (McDonald et al., 2023).

### 2.6 Shotgun metagenome analysis

DNA extracted from the 4th and 12th sponges in each reactor was subjected to short-read metagenomic analysis. Library preparation and sequencing were outsourced to the Bioengineering Lab. Co., Ltd.. In brief, sequencing libraries were prepared using the MGIEasy FS DNA Library Prep Set (MGI Tech, Shenzhen, China) and 150 bp paired-end reads were generated using DNBSEQ-G400 (MGI Tech). Nucleotide sequence data are available in the DDBJ Sequence Read Archive accession number PRJDB18886.

The generated reads were subjected to SOAPnuke 2.0.6 (Chen et al., 2018) to remove the adaptor sequences and low-quality reads. Assembly was performed using the workflow “aviary assemble” in Aviary 0.8.3 (Newell et al., 2023). Contigs shorter than 600 bp were excluded from subsequent analyses. Contigs containing putative *nosZ* gene sequences were retrieved using the BLASTN algorithm (Altschul et al., 1990) and a *nosZ* database (Zheng et al., 2023). The coverage of contigs was analysed coverM 0.7.0 (Aroney et al., 2024), and the relative abundance of each *nosZ* gene was calculated. The obtained contigs were subjected to the workflow “aviary annotate” and Prokka 1.14.6 (Seemann, 2014) for functional annotation. Eventually, contigs containing NosZ protein sequences with ≥ 300 amino acids were extracted to construct a phylogenetic tree.

A phylogenetic tree of the NosZ sequences obtained in this study was constructed using known reference NosZ sequences. The reference sequences were obtained by clustering the *nosZ* gene sequences (Zheng et al., 2023) with 70% sequence identity using VSEARCH 2.27.0. The reference sequences (n=574) were converted to amino acid sequences. The amino acid sequences were aligned using MAFFT (Katoh et al., 2002) and trimmed using TrimAl (Capella-Gutiérrez et al., 2009). A phylogenetic tree was constructed using IQ-tree2 (Minh et al., 2020). The best-fit model was LG + F + R10, as determined using ModelFinder (Kalyaanamoorthy et al., 2017). The phylogenetic tree was visualised using iTOL v6 (Letunic and Bork, 2021).

## 3. Results

### 3.1 N_2_O removal performance by DHS reactors

In this study, the continuous treatment of gaseous N_2_O was conducted using DHS reactors. The N_2_O removal performance was evaluated using different concentrations of N_2_O in N_2_ (5, 40, 300 and 2,000 ppm) under various N_2_O loads by changing the gas flow rates (shortest GRT of 3 min) (Fig. 2 and Tables S1 and S2). Hereafter, each reactor is denoted as 5, 40, 300, and 2,000 ppm N_2_O reactor.

**Fig. 2.**
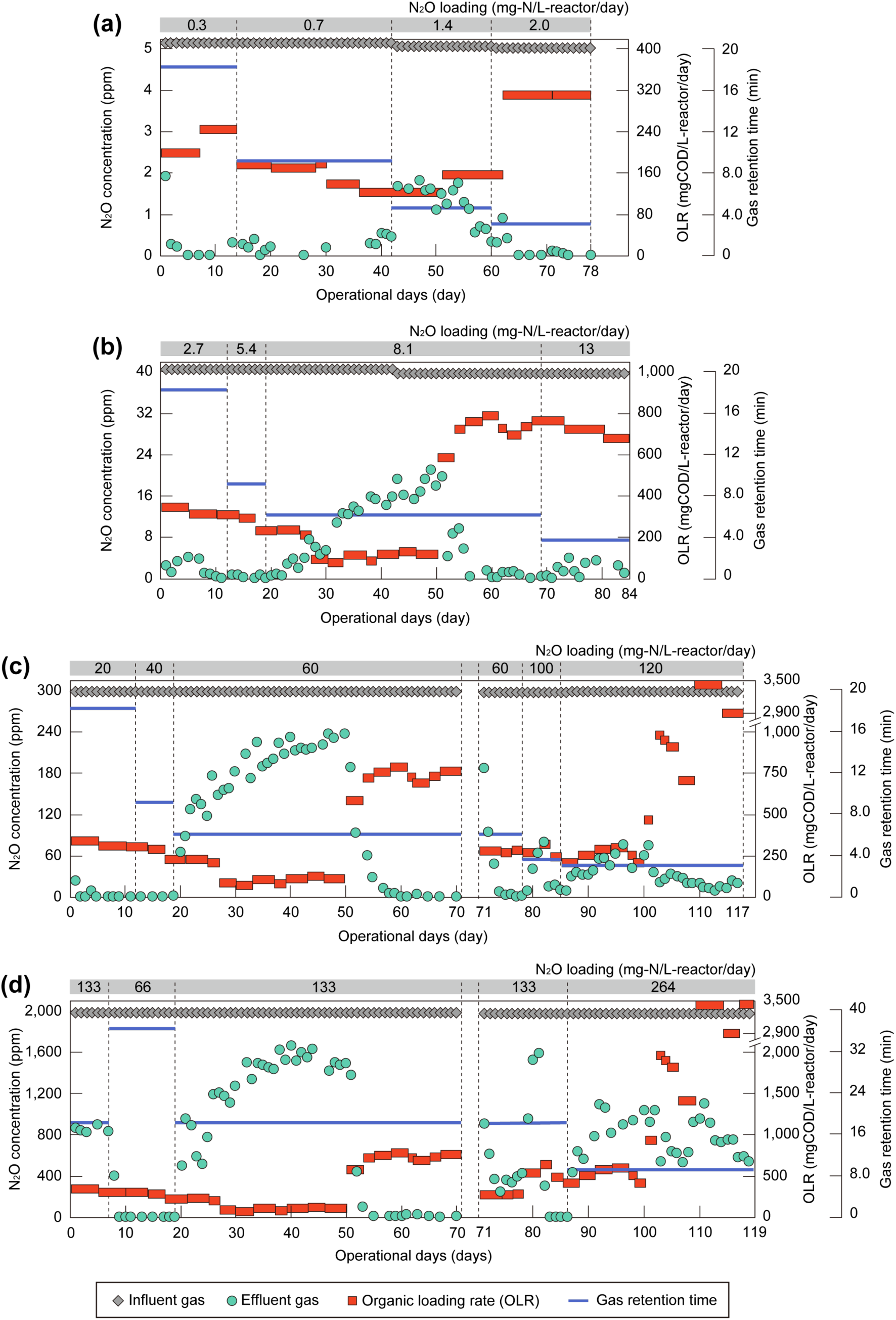
Nitrous oxide (N_2_O) removal performance. Different concentrations of N_2_O in nitrogen gas were treated under different gas retention times and organic loadings. a; 5 ppm N_2_O, b; 40 ppm N_2_O, c; 300 ppm N_2_O, d; 2,000 ppm N_2_O.

The 5 ppm N_2_O reactor (Fig. 2a and Table S1) started with a GRT of 18 min. The N_2_O removal efficiency reached over 90% after one day, and high removal performance was kept even at a GRT of 9.1 and 4.6 min. Finally, the removal efficiency of 97 ± 5.4% was obtained under the GRT of 3.0 min with the N_2_O removal rate of 2.0 ± 0.1 mg-N/L-reactor/day.

The 40 ppm N_2_O reactor (Fig. 2b, Table S1) also showed high removal efficiency after starting up, showing 95 ± 3.9% and 99 ± 0.9% removal efficiency under the GRT of 18 min and 9.1 min by day 20, respectively. When the GRT was shortened to 6.1 min, the removal efficiency dropped to 74 ± 17% (days 20–50) due to insufficient COD supply (Fig. 2b and Tables S1 & S2). After supplying sufficient COD, N_2_O removal became stable with the removal efficiency of 96 ± 3.4% under the GRT of 3.6 min (days 71–85, removal rate; 13 ± 0.5 mg-N/L-reactor/day).

The N_2_O removal performance of the 300 ppm N_2_O reactor (Fig. 2c and Table S1) was similar to the 40 ppm N_2_O reactor. Over 99% removal efficiency was obtained under the GRT as short as 6.1 min (days 61–70), except for the term of days 20–50, during which the COD supply was deficient, and the removal efficiency dropped to 39 ± 16%. On day 70, reactor operation was stopped for unavoidable reasons (Fig. 2c). After a few weeks of suspension, the reactor was restarted with new sponges and seed sludge by feeding 300 ppm N_2_O gas with a GRT of 6.1 min (the day of re-start-up was set as day 71 for convenience). An N_2_O removal rate of over 97% was achieved after four days of operation. The GRT was further shortened to 3.0 min, resulting in the N_2_O removal efficiency of 94 ± 1.5% (days 108–117, removal rate; 113 ± 1.8 mg-N/L-reactor/day).

The 2,000 ppm N_2_O reactor (Fig. 2d and Table S1) was initially operated at a GRT of 18 min. The removal efficiency in the first week was only 57 ± 1.5%; therefore, the GRT was extended to 36 min, resulting in the removal efficiency of 98 ± 6.7%. The GRT was set at 18 min again, and low removal efficiency of 36 ± 18% was observed because of insufficient COD supply (days 20–50) (Fig. 2d and Tables S1 and S2). High removal efficiency of 99 ± 0.6% was achieved with increased organic loading rate (days 61–70). The reactor was stopped after 70 days of operation. Similar to the 300 ppm N_2_O reactor, new seed sludge was inoculated, and the reactor was restarted with a GRT of 18 min. The removal efficiency reached ≥ 99% again after 12 days of operation, and the stable and high removal efficiency of 99 ± 0.1% was observed on days 83–86 (removal rate; 132 ± 0.1 mg-N/L-reactor/day) (Fig. 2d and Table S1). High removal efficiency was not achieved even under higher organic loading rate at the GRT of 9.1 min. Lower N_2_O removal efficiency of 61 ± 10% was obtained (days 87–119), but the highest N_2_O removal rate of 161 ± 26 mg-N/L-reactor/day was achieved in this period.

### 3.2 N_2_O consumption rate

The N_2_O consumption rates of the sludge collected from the 300 ppm and 2,000 ppm N_2_O reactors were measured using microelectrodes, as described earlier (Suenaga et al. 2018) (Fig. S1). Theoretical maximum N_2_O removal rates calculated using the Michaelis-Menten equation were 1,189 ± 257 mg-N/L-reactor/day and 1,468 ± 186 mg-N/L-reactor/day at the 300 ppm and 2,000 ppm N_2_O reactor, respectively (Table 1). Simultaneously, the actual highest N_2_O consumption rates of the reactors were 113 ± 1.8 mg-N/L-reactor/day (days 108–117 for the 300 ppm N_2_O reactor) and 161 ± 26 mg-N/L-reactor/day (days 102–119 for the 2,000 ppm N_2_O reactor). These values were approximately 10% of the theoretical maximum N_2_O consumption rate.

**Table 1.** Nitrous oxide removal rates.

| Reactor | Maximum N<sub>2</sub>O removal rate<br>of the reactor<br>(mg-N/L-reactor/day) | The calculated theoretical N<sub>2</sub>O removal rate<br>(based on the Michaelis–Menten equation)<br>(mg-N/L-reactor/day)<sup>a</sup> |
| --- | --- | --- |
| 300 ppm N<sub>2</sub>O | 113 ± 1.8 (days 108–117) | 1,189 ± 257 |
| 2,000 ppm N<sub>2</sub>O | 161 ± 26 (days 102–119) | 1,468 ± 186 |

<sup>a</sup> N<sub>2</sub>O removal rate per reactor volume was calculated based on the amount of volatile suspended solids retained in the reactor.

### 3.3 Microbial community involved in N_2_O reduction

#### 3.3.1 16S rRNA gene amplicon analysis

The microbial communities of the sludge samples evaluated by 16S rRNA gene amplicon analysis are shown in Table S3, and the top 30 OTUs with average relative abundances are summarised in Fig. 3. The microbial communities built in the reactors differed from those in the seed sludge and substrate, as demonstrated by principal coordinate analysis (PCoA) (Fig. S2). The microbial community structures changed with N_2_O concentrations and along the height of the reactors, although the N_2_O concentrations had the greatest effect.

**Fig. 3.**
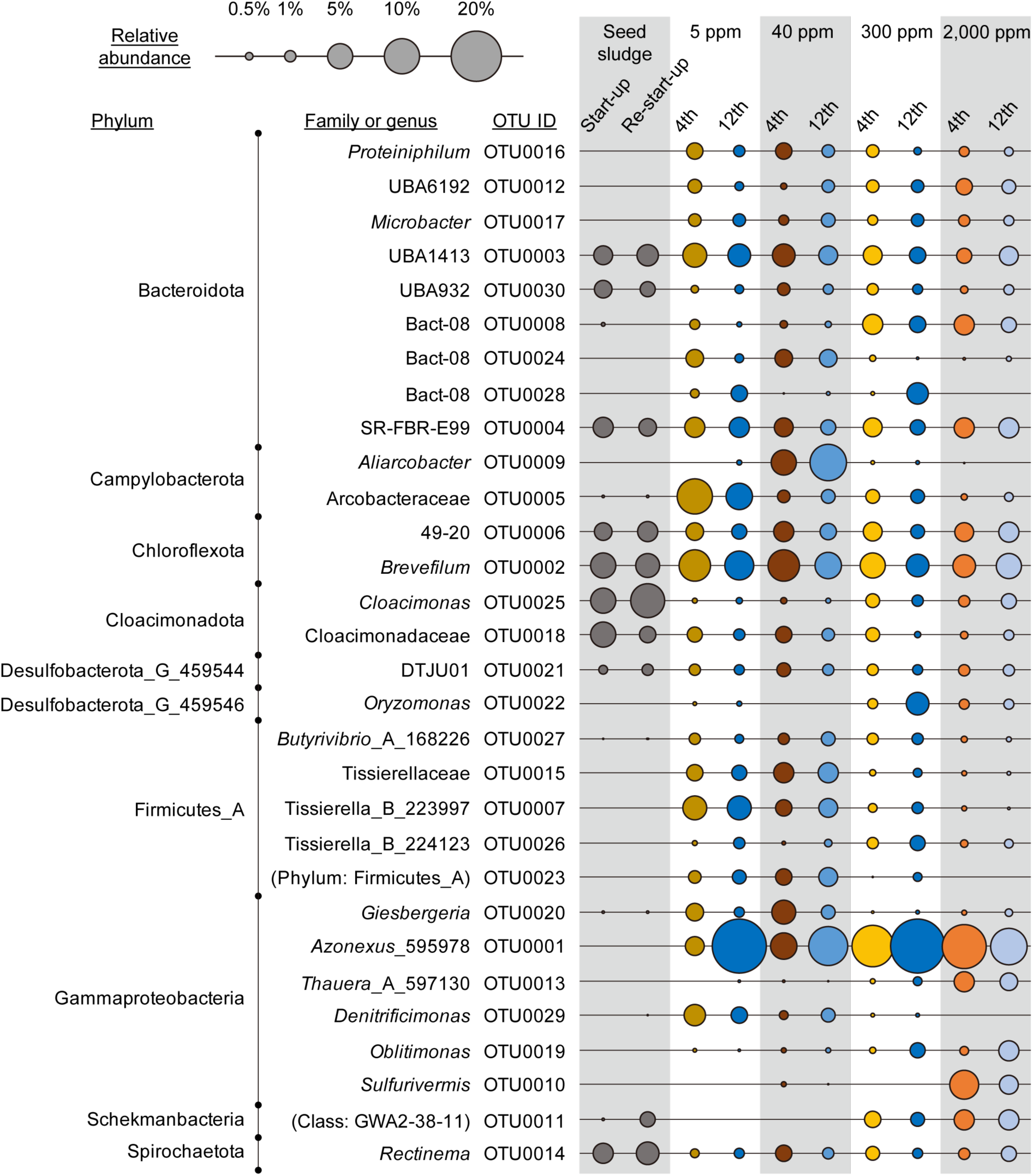
Top 30 OTUs in the average relative abundance. The size of the circle indicates the relative abundance of OTUs in each sample. 4th: the 4th sponge, 12th: the 12th sponge. Seed sludge: sludge from a mesophilic anaerobic sewage sludge digester.

Some OTUs with high relative abundance were similar to known N_2_O-reducing microorganisms (Fig. 3). OTU0001, which is close to *Azonexus*_595978, was the most abundant member in all reactors. *Azonexus* is found in partial-denitrification-anammox processes and anaerobic manure digestate and is expected to be involved in N_2_O reduction (Huang et al., 2024; Wang et al., 2023). In addition to *Azonexus*, several OTUs close to known N_2_O reducers were identified, including *Giesbergeria* (OTU0020) (Zhang et al., 2022), *Thauera*_A_597130 (OTU0013) (Huang et al., 2024), *Denitrificimonas* (OTU0029) (Wang et al., 2024), *Oblitimonas* (OTU0019) (Ramírez-Fernández et al., 2021), and *Sulfurivermis* (OTU0010) (Liu et al., 2023). OTU0020 and OTU0029 were found in reactors with lower N_2_O concentrations of 5 and 40 ppm, respectively, whereas OTU0019 was found in reactors with higher N_2_O concentrations of 300 ppm and 2,000 ppm. OTU0010 was dominant only in the 2,000 ppm N_2_O reactor. Thus, the known N_2_O-reducing microorganisms were enriched in the DHS reactors with various concentrations of N_2_O.

In addition to OTUs close to known N_2_O reducers, the relative abundances of many OTUs increased after feeding with N_2_O. Although they are not close relatives of N_2_O reducers at the genus level, some OTUs belong to the same family as N_2_O reducers (e.g. OTU0005 and OTU0009) (Isokpehi et al., 2024). The OTUs increasing the relative abundance could potentially reduce N_2_O, although this was difficult to demonstrate using only 16S rRNA gene amplicon analysis.

#### 3.3.2 Metagenome analysis of *nosZ* gene

In total, 229 contigs, including a sequence homologous to *nosZ*, were obtained (matched by BLASTN with sequences longer than approximately 600 bp) (Table S4). The phylogenetic tree of *nosZ* extracted from metagenomic 229 contigs (Fig. 4) revealed that clade II *nosZ* gene was more diverse and abundant than clade I *nosZ* gene in the reactors. The most abundant *nosZ* genes retrieved from all reactors were similar to those in *Azonexus*, harboring clade II *nosZ* gene. In the 2,000 ppm N_2_O reactor, other than *Azonexus nosZ* gene, *Sulfurivermis*-related *nosZ* gene was obtained in high abundance. This *nosZ* gene was not detected in the other reactors. Additionally, the *Aliarcobacter*-related *nosZ* gene was only found in the 40 ppm N_2_O reactor, which was identical to the amplicon analysis (i.e. OTU0009). Besides these microorganisms, *nosZ* gene-harboring microorganisms detected in both 16S rRNA gene amplicons and metagenomic analyses were *Giesbergeria* and *Thauera*, both harboring clade I *nosZ* gene.

**Fig. 4.**
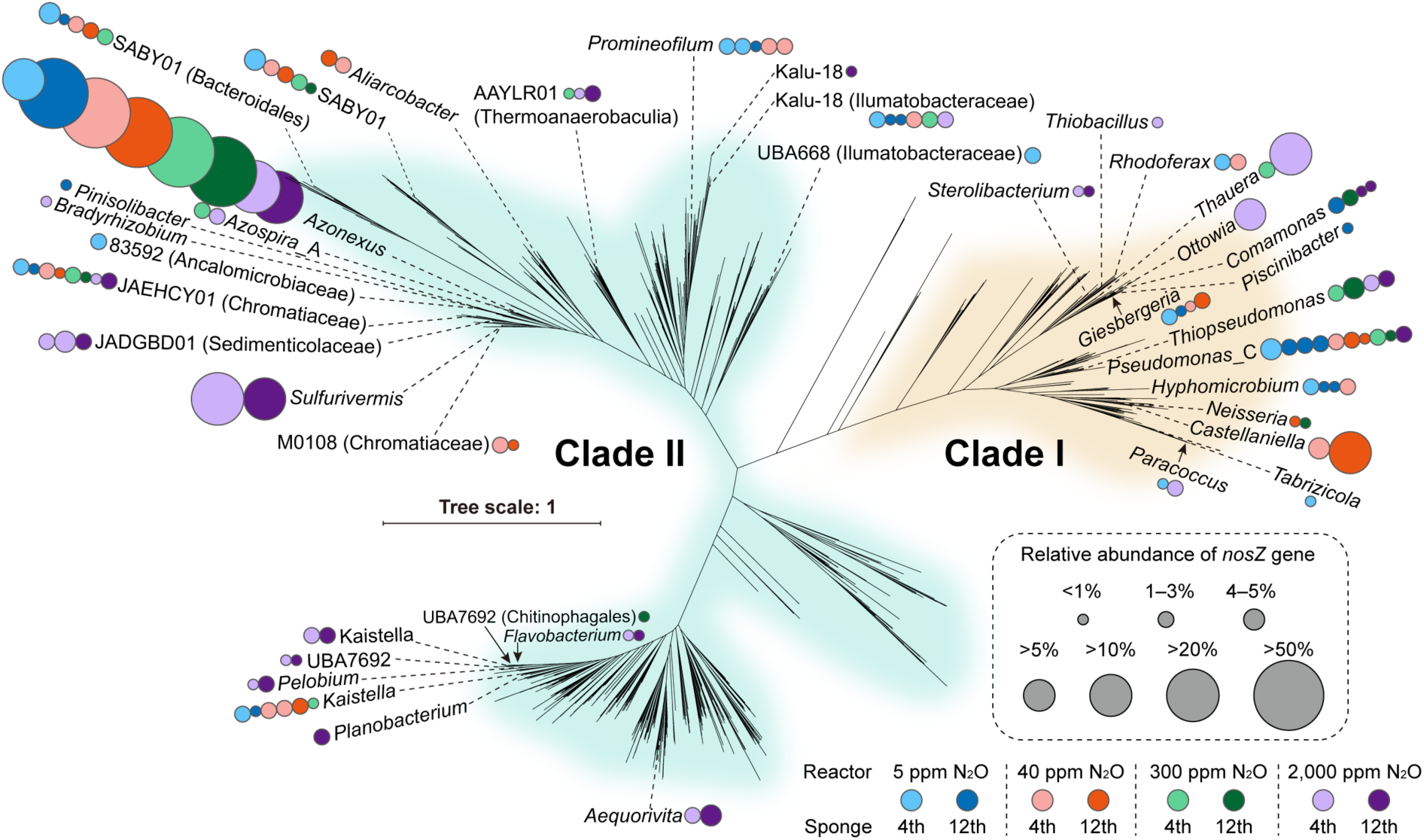
Phylogenetic tree of NosZ protein. The size of the circles indicates the relative abundance of read counts of contigs with the *nosZ* gene. The colours represent the nitrous oxide concentrations fed to the reactors. 4th: the 4th sponge, 12th: the 12th sponge.

## 4. Discussion

### 4.1 Ultra-fast bioprocess for mitigating high concentration N_2_O emission

This study demonstrated that the DHS is able to remove N_2_O in N_2_ gas with removal efficiencies of over 96% for concentrations of up to 300 ppm at 3 min GRT and 2,000 ppm at 18 min GRT with the supply of real wastewater, i.e., the supernatant of sewage sludge digestate. The highest removal rate of 161 ± 26 mg-N/L-reactor/day was achieved when 2,000 ppm N_2_O was supplied with a GRT of 9.1 min. One study investigated the removal of N_2_O in N_2_ gas using a biofilter reactor packed with 2 cm × 2 cm × 2 cm polyurethane cubes (Yoon et al., 2017). This study achieved a removal efficiency of ≥ 99% for 200 ppm N_2_O in N_2_ gas at GRT of 8.9 min with synthetic wastewater and a removal efficiency of 80–90% at GRT of 32.2 min with wastewater from a primary sedimentation basin treating sewage. Our study demonstrated a much faster treatment (over 10 times) capable of removing higher concentrations of N_2_O in N_2_ gas compared to that of a previous study, even with the use of real wastewater.

High concentrations of N_2_O in off-gas were reported, especially from anammox processes, e.g. 0.4–240 ppm (Ali et al., 2016) and 93–1,358 ppm (Okabe et al., 2011) from lab-scale reactors and >4,000 ppm (Kampschreur et al., 2008) from a full-scale reactor although the anammox reaction itself is thought not to produce N_2_O (Harris et al., 2015; Kartal et al., 2007). Although the highest concentration tested in this study (i.e., 2,000 ppm) was lower than 4,000 ppm, it showed the potential to remove such high concentrations of N_2_O from nitrogen gas. A longer GRT is most probably needed, although a higher removal efficiency can be achieved with a sufficient supply of electron donors, which is directly related to the removal efficiencies of the reactor, as shown in Fig. 2bcd (days 20–50 for the 40, 300 and 2,000 ppm reactors).

To minimise the amount of N_2_O emission from wastewater treatment processes, research focusing solely on N_2_O removal efficiency can be misleading. In this study, a removal efficiency of approximately 99% was achieved with a GRT of 18 minutes when 2,000 ppm of N_2_O was supplied. However, approximately 20 ppm of N_2_O still remained in the off-gas. Particularly in cases where high concentrations of N_2_O are treated, implementing an additional treatment process for N_2_O removal from the off-gas can further minimise the final N_2_O emissions from the process (e.g., a series of DHS reactors).

### 4.2 Kinetics of N_2_O reduction in bioreactor

Microelectrode experiments showed an approximately 10-fold faster kinetic potential (Table 1). This suggests that the microbial community inhabiting the reactor had a much greater capacity to reduce N_2_O to N_2_. Dissolved N_2_O was supplied directly to the stirred sludge during the microelectrode experiments (Fig. S1). On the other hand, in the reactor, gaseous N_2_O needs to be initially dissolved in the liquid, and then microorganisms can utilise the dissolved N_2_O. Thus, the N_2_O dissolution rate is the key to determine the removal rate in the reactor.

The dissolution rate from gas to liquid can be explained using the Nernst-Noyes-Whitney equation (Dokoumetzidis and Macheras, 2006; Noyes and Whitney, 1897). In the Nernst-Noyes-Whitney equation, the dissolution rate constant (given by the diffusion coefficient of the substance, volume of the solution, and thickness of the diffusion layer), surface area, concentration of the dissolved substance at a given time t, and saturation solubility of the substance were considered. Without N_2_O consumption in the liquid phase, the dissolution rate decreases as N_2_O dissolves and eventually reaches equilibrium, at which point no more N_2_O is removed from the gas phase. However, when microorganisms are present in the liquid phase and actively consume N_2_O, the N_2_O concentration in the liquid remains low, and the diffusion layer becomes thinner, allowing the dissolution rate to remain high.

To achieve faster dissolution rates, it is suggested to increase the surface area of the carrier or enhance the microbial activity near the sponge surface to decrease the dissolved N_2_O concentration and reduce the thickness of the diffusion layer. One could consider supplying different substrates utilised by other microorganisms to enhance microbial activity to enable faster N_2_O reduction. An earlier study (Yoon et al., 2017) reported a higher N_2_O removal efficiency with synthetic wastewater than that with real wastewater, i.e., effluent from a primary sedimentation basin treating sewage. Additionally, increasing the temperature would help accelerate the microbial N_2_O reduction rate and increase the diffusion coefficient; however, it also causes the decrease of saturation solubility of N_2_O. Decreasing the temperature might help elevate dissolution rates by increasing the saturation solubility of N_2_O although decreased microbial N_2_O reduction rates are expected.

### 4.3 Nitrous oxide reducers in bioreactors

Sludge from the sponges in each reactor was collected on the last day of operation and subjected to 16S rRNA amplicon sequencing analysis. A comparison of the microbial community structure based on PCoA (Fig. S2) showed that the microbial communities in the N_2_O-reducing DHS reactors were different from those in the seed sludge, i.e., sludge from an anaerobic sewage sludge digester, as well as those in the supernatant of an anaerobic sludge digestate, i.e., the substrate for denitrification. It also showed a change in the microbial community under different N_2_O concentrations, as well as in the height of the reactor (Figs. 3 and S2). In a DHS reactor, COD load changes along the height (Kubota et al., 2014), i.e., higher and lower COD concentrations occur in the upper and lower part of the reactor, respectively. Additionally, the supernatant of the anaerobic sludge digestate contains diverse electron donors. These characteristics provide a habitat for a variety of microorganisms in the reactors compared with that of earlier studies using a simple substrate for N_2_O reduction (e.g., acetate for Conthe et al. 2018).

In the N_2_O reducing DHS reactors, *Azonexus* was the most abundant genus after both amplicon and metagenome analyses (Fig. 3 and Table S4). *Azonexus* can utilise intermediates of anaerobic organic degradation such as acetate and propionate (Quan et al., 2006). The digestate supernatant contained a small number of fatty acids. Additionally, the substrate may be anaerobically degraded in the reactor, and the intermediates may be used by *Azonexus* as carbon sources to reduce N_2_O. An earlier study detected *Azonexus* in a bioreactor continuously supplying N_2_O and acetate under anaerobic conditions (Conthe et al., 2018). Thus, *Azonexus* plays a key role in N_2_O removal.

Other than *Azonexus*, *Sulfurivermis*, which can also utilise acetate and propionate for nitrate reduction (Watanabe et al., 2019), was found to be the second most abundant denitrifier, with high relative abundance (26% in the 4th sponge and 17% in the 12th sponge) in the 2,000 ppm reactor. Additionally, *Aliarcobacter* was found in the 40 ppm N_2_O reactor. Because a slightly larger sponge medium (30 mm in diameter and 30 mm in height) was used, a gradient of dissolved N_2_O concentration occurred in the sponge media, which may have involved a variety of microorganisms in N_2_O reduction.

Both clade I and clade II *nosZ* genes were retrieved from the reactors, but clade II *nosZ* was more predominant and diverse than clade I *nosZ* (Fig. 4). *Azonexus*, the most abundant and a key N_2_O reducer of this study, belongs clade II. Generally, microorganisms that possess the clade II *nosZ* gene have lower half-saturation constant (*K_m_*) values and higher affinity than clade I N_2_O-reducing microorganisms and are more abundant in environments where low concentrations of N_2_O are available (Suenaga et al., 2018; Yoon et al., 2016). However, clade II N_2_O-reducers shows higher N_2_O consumption rate than clade I N_2_O-reducers with some specific carbon sources such as acetate and succinate (Qi et al., 2022), suggesting that limited availability of carbon sources derived from the substrate, i.e., the supernatant of anaerobic sludge digestate, caused the predominance of clade II N_2_O-reducers over clade I N_2_O-reducers even under the high concentration of N_2_O.

## 5. Conclusion

In this study, a novel ultra-fast N_2_O removal process using a DHS reactor was developed to remove high concentrations of N₂O generated from anaerobic/anoxic wastewater treatment processes such as an anammox process. Continuous experiments revealed that more than 96% removal rates were achieved for up to 300 ppm N_2_O gas with 3 min GRT and 2,000 ppm N_2_O with 18 min GRT. The maximum N₂O removal rate of 161 ± 26 mg-N/L-reactor/day was obtained when 2,000 ppm N₂O was supplied with the GRT of 9.1 min, which is over 10 times faster than the pioneering process. Additionally, the lack of organic supply immediately deteriorates the N_2_O removal performance. Kinetic analysis indicated that the N₂O dissolution rate is a crucial factor in determining the N₂O removal rate in the reactor. Amplicon and metagenome analyses showed that *Azonexus* played a key role in N₂O removal. A gradient of dissolved N₂O concentration may occur in the sponge carriers, resulting in the involvement of various microorganisms in N₂O reduction.

## Acknowledgements

This study was supported by KAKENHI grants (JP19KK0371, JP21H01460) from the Japan Society for the Promotion of Science (JSPS), the Moonshot project JPNP18016, commissioned by the New Energy and Industrial Technology Development Organization (NEDO) and the MEXT WISE Program for Sustainability in Dynamic Earth (SyDE), Japan. RM was supported by a Grant-in-Aid for JSPS Fellows (JP24KJ0429).

