## Supplemental Figures 1 & 2 for "Mitigating nitrous oxide emission by an ultra-fast bioprocess enabling the removal of high concentration N_2_O"

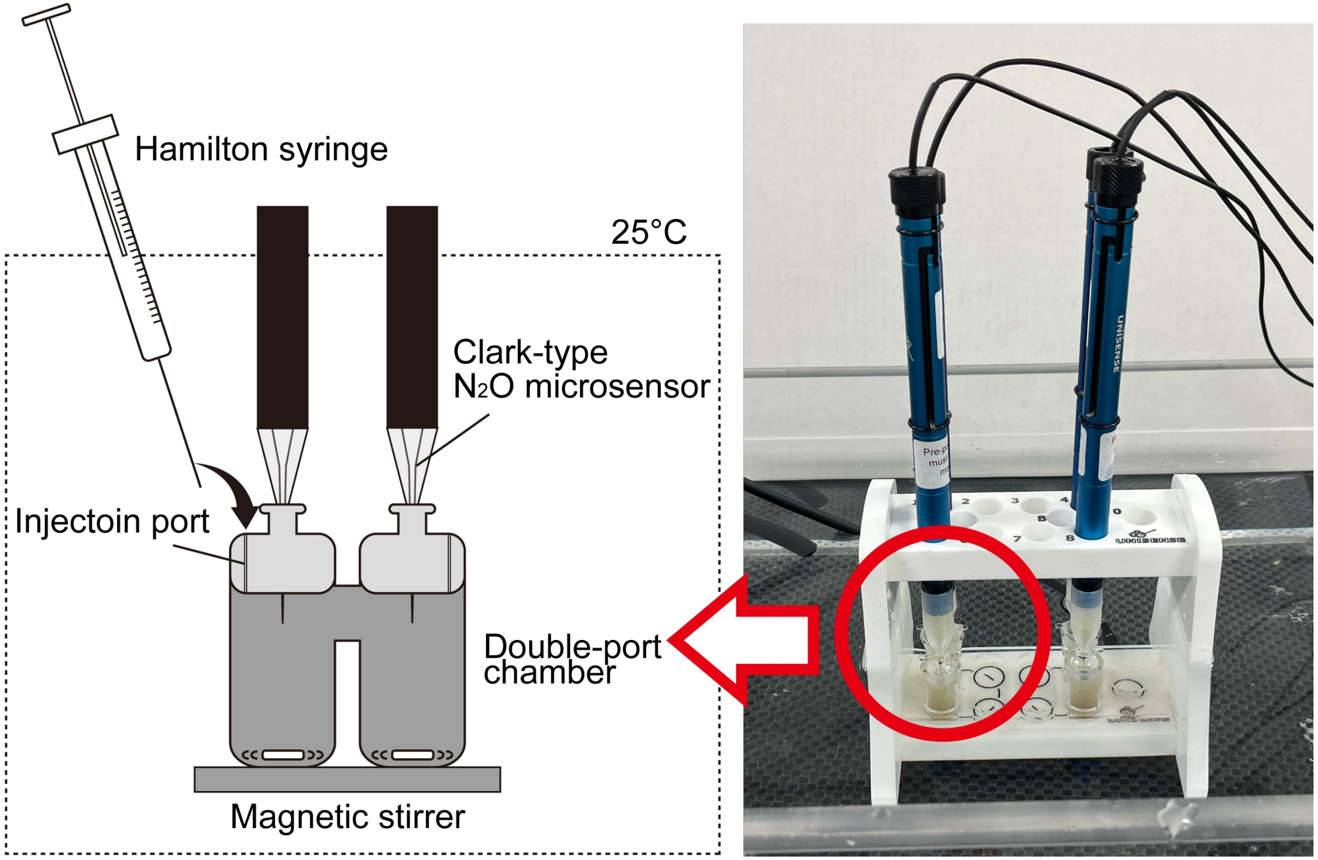


Fig. S1. Diagram of microelectrode experiments.

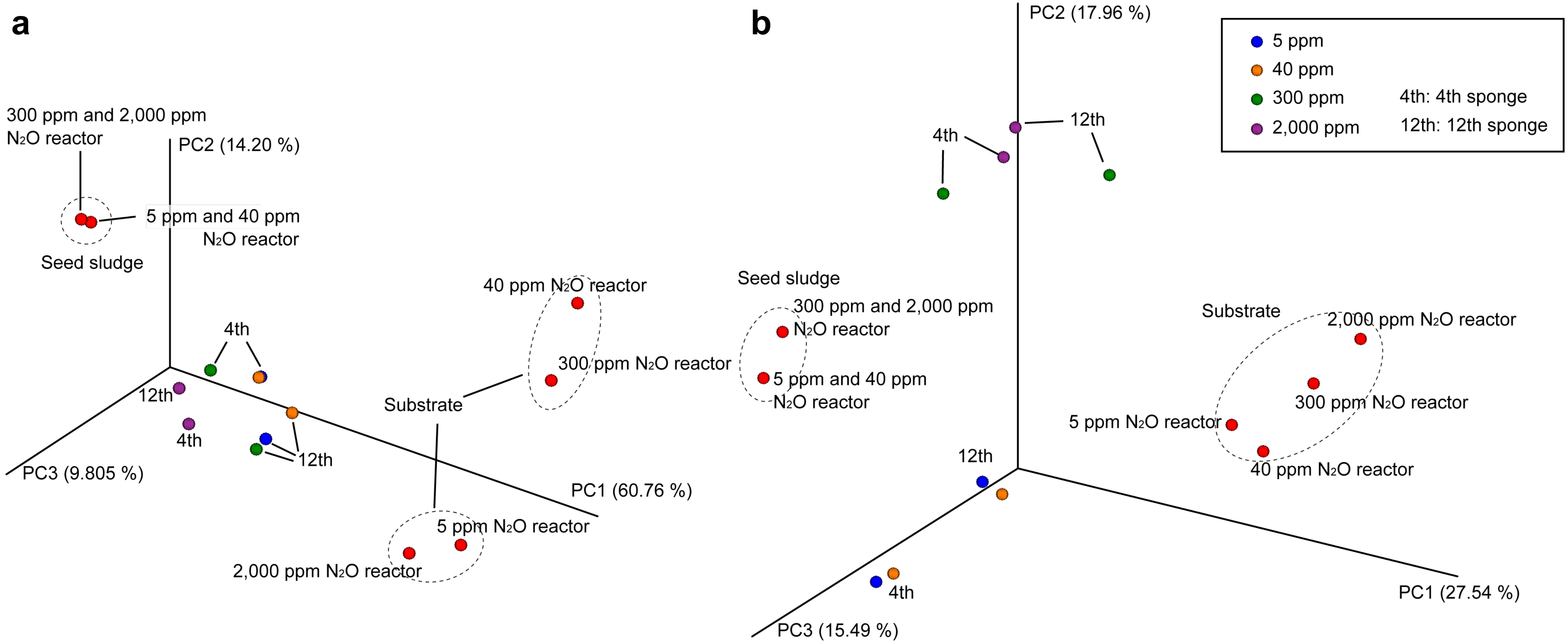


Fig. S2. Principal coordinate analysis with weighted (a) and unweighted (b) UniFrac distance.
4th: the 4th sponge, 12th: the 12th sponge, seed sludge; sludge from a mesophilic anaerobic sewage sludge digester, substrate; supernatant of sewage sludge digester.
